# Conditional Robust Calibration (CRC): a new computational Bayesian methodology for model parameters estimation and identifiability analysis

**DOI:** 10.1101/197400

**Authors:** Fortunato Bianconi, Chiara Antonini, Lorenzo Tomassoni, Paolo Valigi

## Abstract

Computational modeling is a remarkable and common tool to quantitatively describe a biological process. However, most model parameters, such as kinetics parameters, initial conditions and scale factors, are usually unknown because they cannot be directly measured.

Therefore, key issues in Systems Biology are model calibration and identifiability analysis, i.e. estimate parameters from experimental data and assess how well those parameters are determined by the dimension and quality of the data.

Currently in the Systems Biology and Computational Biology communities, the existing methodologies for parameter estimation are divided in two classes: frequentist methods and Bayesian methods. The first ones are based on the optimization of a cost function while the second ones estimate the posterior distribution of model parameters through different sampling techniques.

In this work, we present an innovative Bayesian method, called Conditional Robust Calibration (CRC), for model calibration and identifiability analysis. The algorithm is an iterative procedure based on parameter space sampling and on the definition of multiple objective functions related to each output variables. The method estimates step by step the probability density function (pdf) of parameters conditioned to the experimental measures and it returns as output a subset in the parameter space that best reproduce the dataset.

We apply CRC to six Ordinary Differential Equations (ODE) models with different characteristics and complexity to test its performances compared with profile likelihood (PL) and Approximate Bayesian Computation Sequential Montecarlo (ABC-SMC) approaches. The datasets selected for calibration are time course measurements of different nature: noisy or noiseless, real or in silico.

Compared with PL, our approach finds a more robust solution because parameter identifiability is inferred by conditional pdfs of estimated parameters. Compared with ABC-SMC, we have found a more precise solution with a reduced computational cost.

## Presubmissions for the Methods Section

In Systems Biology, mathematical modeling is a useful activity for understanding many biological processes from a quantitative point of view. It represents an excellent tool to investigate and predict the behavior of some biological species that may not be directly accessible in experimental measurements.

One of the most common modeling techniques consists in representing a biological phenomenon, such as a pathway of biochemical reactions, through a system of ordinary differential equations (ODEs), which describes the dynamic behavior of state variables. Writing an ODE model is a relatively affordable task, using widely known kinetic laws. However, these equations always contain unknown parameters which have to be properly estimated in order to simulate the model and represent the problem under study.

Given specific experimental conditions, the calibration process of a computational model consists in the estimation and inference of parameters in order to make output variables as close as possible to the experimental dataset. Hence, a calibrated model can be used to predict the time evolution of substances for which enough information or measures are not available.

Typically, calibrating a model means identifying those parameters that minimize an objective function, which measures the error between simulated and experimental data.

Moreover, it is also significant to assess how much uncertainty there is in the parameter estimate returned by the identification processes. To this purpose, identifiability analysis is usually combined with parameter estimation and can be performed through an a priori or data-based approach, whether or not experimental data are available [1].

The most common methodologies for parameter estimation and identifiability analysis can be divided in two classes: the frequentist approach and the Bayesian approach [2] [3].

The frequentist methods aim at maximizing the likelihood function, which is the probability density of observing the dataset given certain parameter values. These techniques implement a global and/or local optimization algorithm and return as output a solution that is the best fit between simulated and real data.

Furthermore, for this category of approaches, identifiability analysis corresponds to the estimation of confidence intervals for all model parameters. One of the most widely used approaches in Systems Biology and Computational Biology is the so-called Profile Likelihood, a data-based algorithm that is able to distinguish structural and practical non-identifiability. Profile likelihood starts from the best fit and, for each parameter, subsequently changes its value of a fixed step and then re-optimizes all the others in order to obtain the confidence interval of the given parameter [4].

Parameters are declared as non-identifiable if they exhibit an infinite confidence interval, which means that the parameter itself does not affect the behavior of output variables. The main drawbacks of this method are the possible existence of local optima, the overfitting of data and the lack of robustness of estimated parameters [5]. Since confidence intervals are independent among each other, perturbing one parameter within its confidence region does not imply that all the others range in their corresponding confidence intervals. Thus, PL does not identify a joint region in the parameter space that guarantees the desired behavior for all output variables.

On the other hand, the Bayesian approach considers parameters as random variables, whose joint posterior distribution is estimated through the Bayes theorem. More in detail, the posterior distribution is the conditional probability density function (pdf) of parameters given an experimental dataset. The joint posterior density automatically provides an indication of the uncertainty of the parameter inference and gives major insights about the robustness of the solution. Thus, these algorithms seek to identify regions of high probability density in the parameter space.

Since computing the posterior distribution analytically is usually not feasible, sampling based techniques are used to estimate it. Two classes of sampling methods widely used are the Markov chain Monte Carlo (MCMC) and the Sequential Monte Carlo (SMC). MCMC algorithms approximate the posterior distribution with a Markov chain, whose states are samples from the parameter space. The SMC algorithms evaluate an approximation of the posterior distribution through a series of intermediate distributions, obtained by iteratively perturbing the parameter space. These methods define a single distance function between simulated and experimental data and a series of predefined thresholds. Each iteration then selects only those parameters that give rise to a distance function under the threshold. The main limits of these methods are the computational burden and the curse of dimensionality [6].

The purpose of this paper is to introduce a novel computational approach, called Conditional Robust Calibration (CRC), for parameter estimation of mathematical models, in order to identify a region in the parameter space yielding a specific behavior. The proposed method is an iterative procedure whose mainstays are the parameter space sampling and the estimation of the probability density function (pdf) for each model parameter. For these reasons, CRC is located in the category of the Bayesian algorithms. Despite that, it presents many novel aspects and features that makes it unique compared to all the other existent methodologies. Indeed, it is important to highlight that we did not apply, extend or modify any of the algorithms published in the literature.

Our algorithm consists of the following steps: once a distance function is defined for each output variable, at each iteration, we perturb the parameter space with a fixed number of samples, we compute the distance functions and we estimate their probability density functions after parameter perturbation, through a kernel density approach. At each step, we accept only those parameter values for which each distance function is under a specific threshold. Each threshold is chosen in order to guarantee the convergence of the conditional parameter density functions to a steady state, given the region of interest. Since the purpose is to shift each threshold toward zero, as the number of iterations increases, the range of parameter perturbations is progressively tightened so that samples are thickened near the possible solution. This process is repeated until the desired level of agreement between the simulations and the dataset is reached. In the end, the algorithm returns a robust solution because it selects a region in the parameter space that gives rise to the desired behavior. The pdf of each parameter in the last iteration gives information about the identifiability and uncertainty of the parameter.

One of the main innovations in our methodology is the overturning of the sampling approach: the number of samples is unchanged throughout iterations and must be fixed initially by the user. This innovation is substantial because, from the beginning, there is complete knowledge and control of the computational costs and thus it is possible to accurately estimate how long the iteration will take. This sampling strategy guarantees that the computational time does not increase. It actually sometimes diminishes as perturbation intervals are shrunk. On the other hand, in SMC, the input value is the threshold to reach at each iteration and thus the required number of samples is not known a priori. As a result, since the threshold decreases at each step, the number of generated samples could increase as far as the time of the computational simulation.

In order to reach a uniform coverage of the parameter space, we employed the Latin Hypercube Sampling (LHS) technique because it allows a joint perturbation of all parameters. Another innovation is the comparison of the performances of our algorithm using logarithmic and linear LHS in each validation example. This allowed us to infer the importance of the sampling approach and to discover the best sampling technique between the two types described above for a correct calibration of a model.

Another distinct characteristic of this new method is the definition of as many objective functions as output variables, with the aim of minimizing all of them, in order to provide a more balanced and precise calibration toward each measure of the dataset. This is relevant because, on the contrary, both the Bayesian and the frequentist class minimizes a single objective function for all observables. Consequently, there is no information about the error on the single output variable.

Finally, the last innovation concerns identifiability analysis of model parameters. Indeed, in addition to the evaluation of the conditional parameter densities, for each parameter we computed the Moment Independent Robustness Indicator (MIRI), which is an index that measures the dissimilarity between two pdfs [7]. In addition to the pdfs in output from CRC, we also estimated the conditional pdfs of all parameters, given high values of the distance functions. Then, for each parameter, we computed the level of intersection between the two pdfs defined above, through the MIRI indicator. In this way, it is possible to quantify the influence of each parameter on the behavior of interest. Parameters with higher values of the MIRI have a major impact on output variables because there is a larger shift between the two conditional densities used for MIRI calculation.

We validated this new methodology on six different ODE models of increasing complexity and each one with specific features, in order to demonstrate the flexibility of our approach. All the models are already published in Systems Biology journals and they combine the most widely used kinetic laws.

In all the examples, we also applied the Profile Likelihood Approach, through the software Data2Dynamics, and the Approximate Bayesian Computation (ABC) SMC, through the ABC-SysBio software [8], with the purpose of having a reliable and complete comparison with the state of the art of this field.

The first three models are toy models that describe the basic common kinetic laws: the exponential decay, Michaelis Menten kinetic and the law of mass action. They have, respectively, one kinetic parameter and one output variable, two kinetic parameters and one output variable, three kinetic parameters and two output variables. All of them were calibrated using an in silico noiseless dataset.

The fourth case study is the Lotka-Volterra model, consisting of two equations and two kinetic parameters, with the distinctive feature of an oscillatory behavior of both output variables. It was calibrated on an in silico noisy dataset [6].

The fifth example represents the activation of an enzyme via two steps and it has seven parameters and two output variables [9]. Differently from the previous cases, parameters to estimate are not only kinetics but also initial conditions and scale factors. It was calibrated on a simulated noisy dataset.

The last example is an ODE model of the signalling pathway of p38 MAPK in multiple myeloma (MM). It has 16 output variables and 53 kinetic parameters and it was calibrated on a Reverse Phase Protein Array (RPPA) experimental dataset [10].

CRC was successful in all cases, especially when we applied the logarithmic LHS as sampling technique. Compared to the profile likelihood approach, we always obtained a more robust solution because confidence intervals of parameters are described through marginal distributions of the joint posterior density of the whole parameter vector.

On the other hand, compared to the Bayesian approach, our method reached a more precise solution with a definitely lower computational cost, mainly because we defined multiple objective functions and a fixed number of parameter samples.

All these advantages are evident in all the models studied (see Figure 1 and 2 as result example) but especially in the most complex model, showing that our procedure can be applied to models with a parameter space of high dimension and a dataset resulting from an high-throughput experiment. The ABC-SysBio software applied to this model after 10 days of simulation did not finish. On the other hand, the Profile Likelihood Approach estimated degenerate confidence intervals for most parameters, i.e., confidence intervals where upper and lower bound coincide. Thus, for these parameters, it is not possible to assess their identifiability.

**Figure 1:**
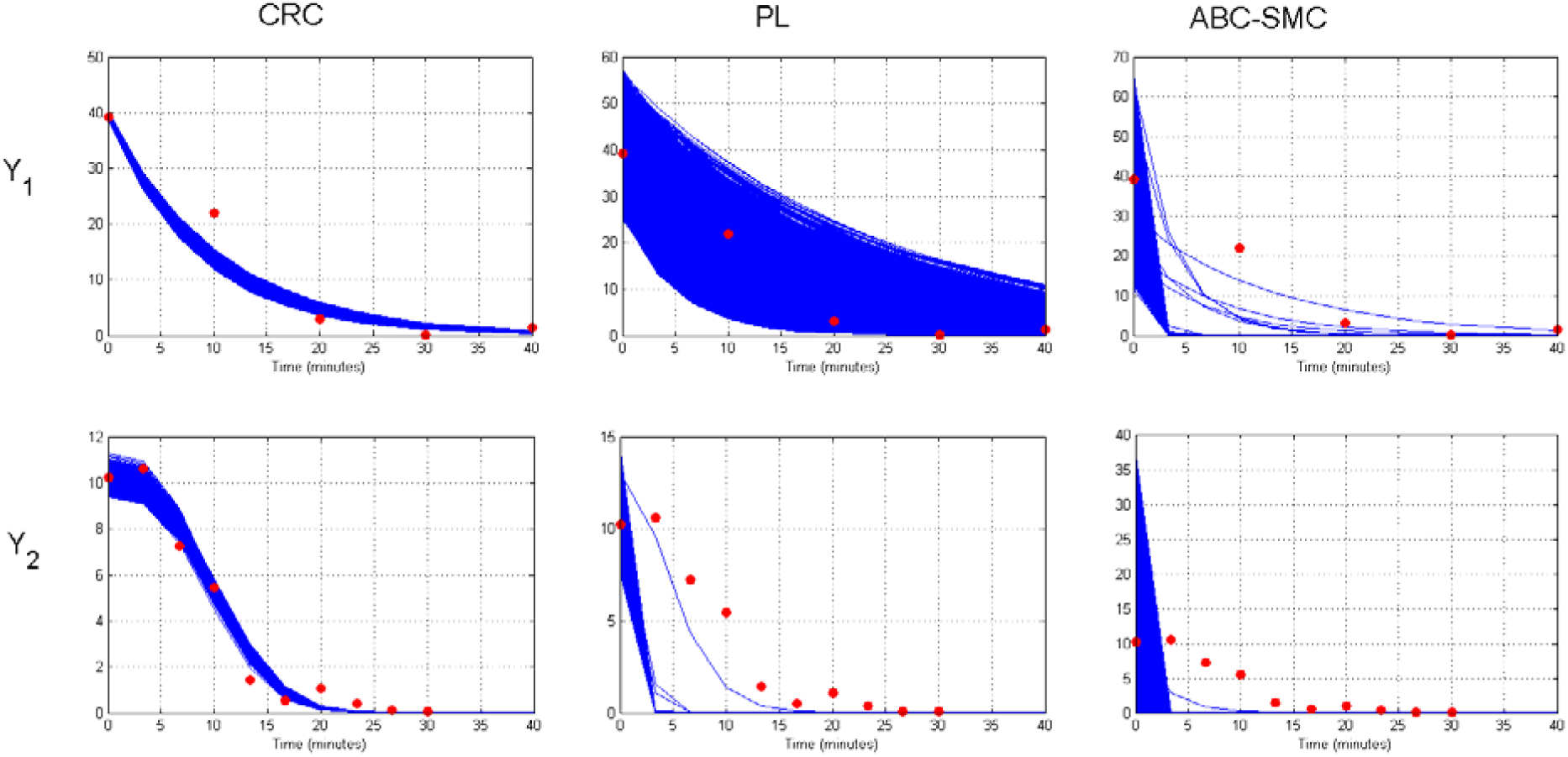
Results of the three calibration methodologies applied to the ODE model in [9]. Time behavior of output variables Y_1_ and Y_2_ compared between CRC, PL and ABC-SMC. Blue lines are the time simulations and red dots represent the in silico noisy dataset used for calibration. (a) Simulations were performed using parameter values belonging to the region obtained in the last iteration of CRC. (b) Robustness of the profile likelihood solution: parameters are simultaneously perturbed in their corresponding confidence intervals returned by PL. (c) Parameters are set equal to the particles returned by ABC-SMC after computation of the last parameter population.

**Figure 2:**
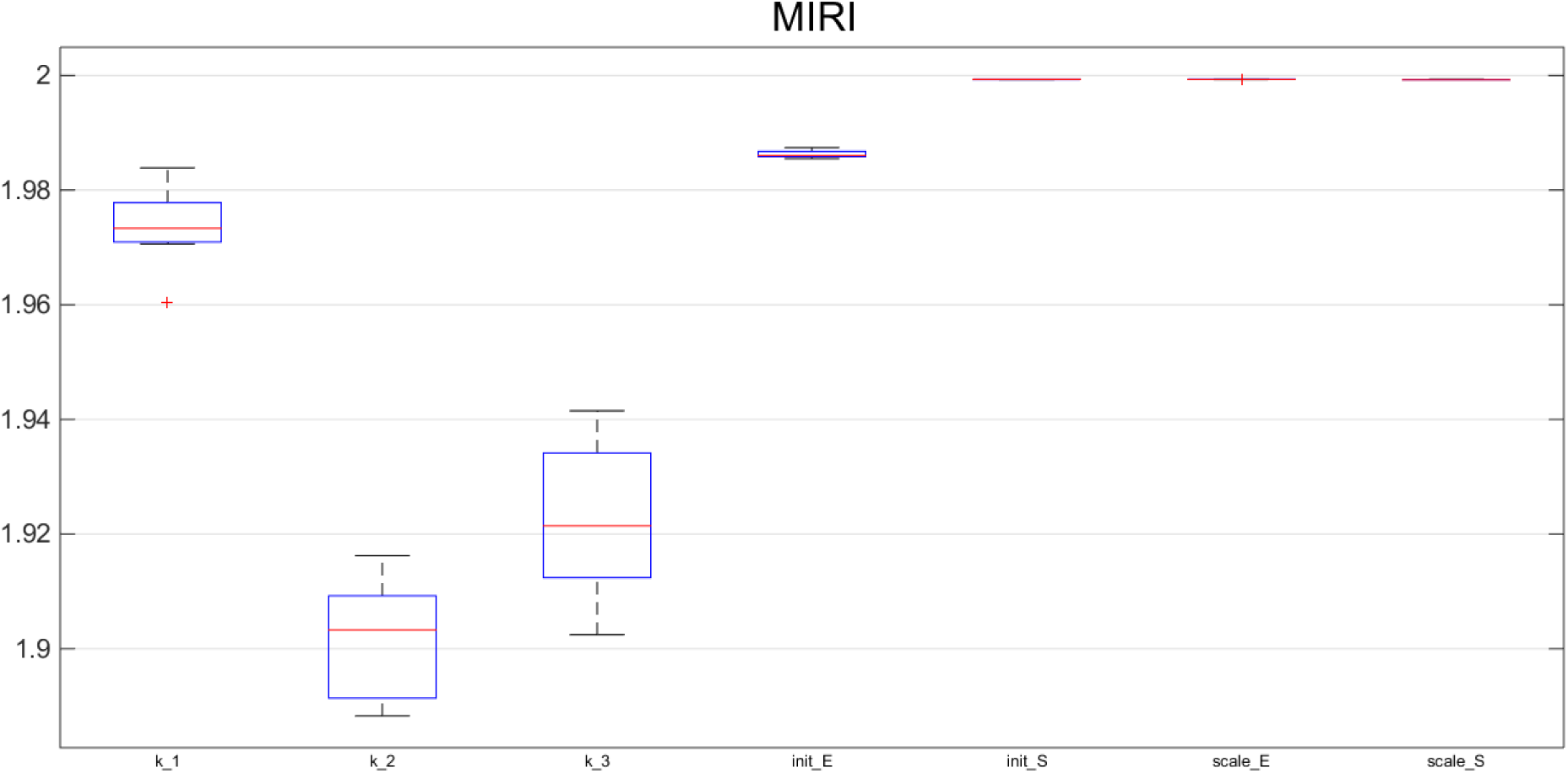
MIRI for model parameters of [9]. These results are in agreement with those of the PL in [9]. Both k_2_ and k_3_, which are non-identifiable according to the PL algorithm, have MIRI lower than those of the other parameters of the model.

We developed CRC as a set of Matlab functions, whose fundamental source code is freely available at <u>CRC github project.</u>

## References

1: Villaverde AF, Barreiro A, Papachristodoulou A. Structural Identifiability of Dynamic Systems Biology Models.Plos Computational Biology.2016; 12 (10):e1005153

2: Lillacci G, Khammash M. Parameter Estimation and Model Selection in Computational Biology.Plos Computational Biology.2010; 6 (3):e1000696

3: Frohlich F, Kaltenbacher B, Theis FJ, Hasenauer J. Scalable Parameter Estimation for Genome-Scale Biochemical Reaction Networks.Plos Computational Biology.2017; 13(1)::e1005331

4: Raue, A., Kreutz, C., Maiwald, T., Bachmann, J., Schilling, M., Klingmüller, U., & Timmer, J. Structural and practical identifiability analysis of partially observed dynamical models by exploiting the profile likelihood.Bioinformatics.2009; 25 (15):1923–1929

5: Sontag ED. Dynamic compensation, parameter identifiability, and equivariances.Plos Computational Biology.2017; 13 (4):e1005447

6: Toni, T., Welch, D., Strelkowa, N., Ipsen, A., & Stumpf, M. P. Approximate Bayesian computation scheme for parameter inference and model selection in dynamical systems. Journal of the Royal Society Interface.2009; 6 (31):187–202

7: Bianconi, F., Baldelli, E., Luovini, V., Petricoin, E. F., Crinò, L., & Valigi, P. Conditional robustness analysis for fragility discovery and target identification in biochemical networks and in cancer systems biology. BMC systems biology.2015; 9 (1):70

8: Liepe, J., Kirk, P., Filippi, S., Toni, T., Barnes, C. P., & Stumpf, M. P. A framework for parameter estimation and model selection from experimental data in systems biology using approximate Bayesian computation. Nature protocols.2014; 9 (2):439–456

9: Raue, A., Kreutz, C., Maiwald, T., KlingMüller, U., & Timmer, J. Addressing parameter identifiability by model-based experimentation. IET systems biology.2011; 5 (2):120–130

10: Peng, H., Peng, T., Wen, J., Engler, D. A., Matsunami, R. K., Su, J., … & Zhou, X. Characterization of p38 MAPK isoforms for drug resistance study using systems biology approach.Bioinformatics.2014; 30 (13):1899–1907

